# The evolutionary dynamics of plant mating systems: how bias for studying ‘interesting’ plant reproductive systems could backfire

**DOI:** 10.1101/2024.06.18.599380

**Authors:** Elena M. Meyer, Laura F. Galloway, Andrew J. Eckert

## Abstract

**Background and Aims:** An “abominable mystery”: angiosperm sexual systems have been a source of both interest and frustration for the botanical community since Darwin. The evolutionary stability, overall frequency, and distribution of self-fertilization and mixed-mating systems have been explored in a variety of studies. However, there has been no recent study which directly addresses our knowledge of mating systems across families, the adequacy of existing data, or the potential for biases.

**Scope:** Here we present an updated dataset of mating systems across flowering plants covering 6,781 species and 212 families based on a synthesis of existing reviews and an original literature review using Web of Science. We assess the adequacy of this data by evaluating for bias indicating enrichment of certain families or sexual systems.

**Key Results:** We find that the vast majority of our data on mating systems comes from a small number of disproportionally sampled families, and that families with significant proportions of dioecious or monoecious species are much more likely to be undersampled.

**Conclusions:** Our results show that the frequency of selfing in angiosperms is overestimated, possibly due to increased research interest in selfing and mixed-mating systems. This suggests that systematic study bias may mean we know less about this vital facet of plant life than we think.

## Introduction

Darwin famously described the rapid emergence and evolutionary radiation of angiosperms as an “abominable mystery” (Darwin 1876, compiled 1903; Friedman 2009). One key to this mystery lies in the diverse reproductive strategies possessed by angiosperms. Reproduction is a key determinant of fitness (Charlesworth 1994; Kalisz 1989), and thus it is a fundamental evolutionary driver. Despite this importance, our present understanding of the frequency and distribution of different reproductive strategies in angiosperms remains both limited and fraught with the potential for bias (Igić and Kohn 2006).

Plants possess an unusual level of variation in their reproductive strategies (Harder and Barrett 2006). Within angiosperms, reproduction can take place either asexually (via apomixis or by vegetative clonal growth) or sexually (2006). For sexual reproduction, two modes are possible: either self-fertilization (“selfing”) or outcrossing involving two parental plants. Many plant species, and even the same individuals within a species, perform both strategies, known as mixed mating (Stebbins 1950; Goodwille et al. 2005). In primarily outcrossing species, mechanisms often exist to avoid reproduction through selfing (e.g., genetic incompatibility with self pollen, anatomical separation of male and female flowers, or divergent flowering time) (Lande and Schemske 1985). The diversity of mechanisms that plants use to prevent selfing suggests that outcrossing is often advantageous. Avoidance of selfing has been associated with higher family diversification rates (Ferrer and Good 2012), providing some evidence for this theory. However, the known phylogenetic breadth of selfing implies that despite these avoidance mechanisms, conditions exist where selfing is favored, at least in the short-term.

Designation of sex in plants is also varied: individuals may be dioecious, with separate sexes corresponding to individuals; monoecious, with separate sexes corresponding to flowers on a single individual; or hermaphroditic, with both sexual organs present in the same floral structure. Self-fertilization is impossible for dioecious species due to their morphology. While selfing is possible for both monoecious and hermaphrodite species, in monoecy pollen must travel from the male to female flower for selfing to occur (i.e., geitonogamy). This requires either wind or pollinator facilitation. Dioecy and monoecy present an additional opportunity to enforce outcrossing through spatial or temporal separation of the sexes. Conversely, hermaphroditism could encourage selfing through proximity of the male and female parts of the flower. In this way, the ease of selfing for different sexual systems varies along a spectrum: selfing is impossible for dioecious species, more challenging for monoecious species, and most likely for hermaphroditic species. Because of this correspondence, studies of plant sexual systems also provide information about mating systems. Mating systems and sexual systems are often addressed separately (for example, Goodwille et al. 2005 treats only mating systems, Wang et al. 2021 treats only sexual systems; but see Holsinger 2000). However, knowledge about sexual systems clearly informs knowledge about mating systems. Both are key to the unrivaled diversity of angiosperms (Barrett 2002), suggesting that additional synthesis of these traits will be beneficial to our understanding of reproduction in plants.

Considerable theoretical work has evaluated the evolutionary costs and benefits of selfing (Darwin 1876; Fisher 1941; Nagylacki 1976; Lloyd 1977, 1979, 1980; Cheptou 2019). Most straightforwardly, self-fertilization holds a transmission advantage of gene copies over two-parent reproduction (Fisher 1941). Based on this alone, without an evolutionary cost to selfing, it would become commonplace. The obvious evolutionary cost to selfing is the increased expression of recessive deleterious phenotypes leading to inbreeding depression. Thus, to explain the variation we observe in plant reproductive systems, we must explain both the continued existence of selfing plants in the face of inbreeding depression and the persistence of outcrossing in spite of the two-fold transmission advantage of selfing. The fitness cost incurred by inbreeding depression (Charlesworth and Charlesworth 1987) may explain why selfing does not outcompete outcrossing based on transmission advantage alone (Lloyd 1979). Under circumstances where the cost of inbreeding depression does not outweigh the transmission advantage of selfing, however, selfing can persist (Lloyd 1979). Another commonly cited advantage of selfing is reproductive assurance in the absence of a mate, or pollen limitation (Darwin 1876; Kalisz 2004; reviewed by Busch and Delph 2012). Selfing has thus also been proposed as a mechanism to aid colonization of new habitats, with self-compatible organisms more likely to be successful colonizers than self-incompatible species (Baker 1955). This is also sometimes referred to as “Baker’s Law.”

From this body of literature, two prominent – but not mutually exclusive – hypotheses emerged about the expected frequency of selfing across angiosperms. Stebbins (1957) proposed that selfing is an evolutionary “dead end,” with selfing species more prone to extinction. He suggested that the limitations placed on genetic diversity by selfing would constrain adaptation to new environments. This hypothesis suggests, even if the deleterious alleles associated with inbreeding depression could be purged, that over time the loss of genetic diversity would doom selfing species to extinction. However, if transitions to selfing were sufficiently frequent, at any given point in evolutionary time, we could still potentially observe extant selfing species (Stebbins 1957).

Schemske and Lande (1985) proposed that the distribution of selfing should be bimodal, with most species either primarily selfing or primarily outcrossing. Outcrossing would represent a stable strategy by avoiding inbreeding depression and promoting diversity through biparental sexual reproduction. Similarly, highly selfing species would be able to avoid inbreeding depression by purging deleterious alleles and would thus persist due to their transmission advantage and reproductive assurance. Species with mixed mating, however, should be less common, as purging of deleterious alleles would be more difficult under intermediate levels of selfing, yet the risks of inbreeding depression and loss of genetic diversity remain.

Since Schemske and Lande (1985), the distribution of selfing across seed plants has been revisited multiple times. Vogler and Kalisz (2001) found that the bimodal distribution hypothesis was supported for wind-pollinated species, but not for animal-pollinated species. This was revised again when Goodwille et al. (2005) reviewed mixed mating (defined in their review by an outcrossing rate, *t,* falling between 0.2 and 0.8). They showed that mixed mating systems are more frequent than previously thought (in their study, up to 42% of species selfed or had the potential to self), and the frequency at which mixed mating is observed to occur suggested some level of stability for selfing. The authors additionally cautioned, however, that their data was “far from random,” and drew 44% of its total species from only five families (Goodwillie et al. 2005). Shortly after, Igić and Kohn (2006) demonstrated a bias against obligate outcrossing species in published studies of outcrossing and selfing rates and cautioned that any generalizations about the true distribution of selfing were likely premature.

All these studies (Schemske and Lande 1985, Vogler and Kalisz 2001, Goodwille et al. 2005) explicitly address the question of the true underlying distribution of selfing. Since the publication of these seminal papers, the rich and provocative theoretical background surrounding selfing has promoted a wealth of empirical papers addressing selfing in various plant systems (for many good examples, see Whitehead et al. 2018). This has greatly expanded the dataset of plant species with known mating systems. Schemske and Lande (1985) presented a dataset of just 55 species to demonstrate the bimodal distribution empirically. Now, known mating systems number in the thousands, as shown in this review. However, this larger dataset has the potential to continue promoting the existing biases noted by Igić and Kohn (2006).

Existing studies fall into one of three categories: (1) papers that directly address selfing rate evolution and selfing frequency across multiple plant families, (2) papers that address the interplay of the selfing trait with other topics in ecology and evolution, or (3) studies that focus on specific families that are known to have varied or potentially labile mating systems. A focus on a few families with especially varied mating systems is often necessary for individual studies. But over time, this will inevitably lead to nonrandom and biased sampling across the angiosperm phylogeny with respect to mating systems, since the focus is on selfing species and not producing random samples across the angiosperm phylogeny (*cf*. Igić and Kohn 2006).

Papers that address the interplay of selfing at a multi-family level with other various topics in ecology and evolution have been especially prolific. For example, selfing has been explored in conjunction with geographic range dynamics (Grossenbacher et al. 2015, 2016), invasive status and naturalization (Rananjatovo et al. 2016), island biogeography (Grossenbacher 2017), within-population selfing rate variation (Whitehead et al. 2018), life history and range size (Prior and Busch 2021), and seed size (Tateyama et al. 2021). Additionally, studies focused on estimating rates of transitions to and from selfing using phylogenetic comparative methods (PCMs) have contributed significantly to knowledge about the tempo and mode of selfing evolution. Such studies have provided detailed knowledge about the role of selfing to diversification rates in Solanaceae (Goldberg et al. 2015), Onagraceae (Freyman and Hoha 2019), and Fabaceae (Delaney and Igić 2021). These families were selected because they have labile mating systems. While this is appropriate for the goals set forth in each study, this reinforces existing biases against families with high frequencies of outcrossing. Thus, our collective knowledge about the underlying distribution of selfing, if based on meta-analyses using these studies, will likely be biased.

Numerous reviews focusing more broadly on selfing also exist (e.g., Goodwille et al. 2005, Wright et al. 2013). However, the question of the underlying distribution of self-compatibility (and, conversely, self-sterility) has not been recently revisited. Therefore, we remain extremely uncertain about what percentage of angiosperm species are capable of selfing. This means we also don’t know if our data allow us to estimate trends across angiosperm-wide diversity. In previous work, the significant proportion of known mating systems that incorporate selfing has been used to suggest the stability of mixed mating (Goodwille et al. 2005). However, if selfing and mixed mating are selectively over-sampled, this assumption could be premature. Similarly, if our understanding of the true underlying distribution of selfing is biased, then we run the risk of making incorrect and overly simplifying assumptions when modeling mating system evolution with use of PCMs. Incorrect or biased assumptions, moreover, may more generally lead us to incorrect conclusions, even when estimating quantities as simple as the frequency of selfing and mixed mating across angiosperms.

The evolutionary dynamics of plant mating systems are more than just a challenging theoretical problem. They also quantify fundamental driving forces determining fitness in real plant populations. Allocation of selfing and outcrossing across individuals varies widely within populations and is labile in populations responding to environmental change (Whitehead et al. 2018). Theory suggests that selfing may be adaptive in rapidly changing environments by providing reproductive assurance, especially in situations of pollinator limitations (Darwin 1876, Baker 1955, Cheptou 2019). However, selfing may also limit adaptive potential, especially in the long term (Stebbins 1957). In an era of rapid anthropogenic climate change, some species have already changed their reproductive allocation in response to climate (Rech 2021; Austin 2022, Cheptou et al. 2022). Similarly, significant pollinator decline has been a hallmark of our era (Thomman et al. 2013) and can create pollen limitation (Ashman et al. 2004). This could lead to evolutionary pressures promoting the rapid change of reproductive traits to adapt (Thomman et al. 2013). Thus, in the Anthropocene, selfing could be favored in the short-term by climate instability but lead to extinction in the longer term as the known costs to selfing are realized. However, some species may have stress-induced mechanisms to promote outcrossing as opposed to selfing during episodes of drastic environmental change (van Ginkel and Flipphi 2020). It is thus paramount to better understand the existing biases in our knowledge about the underlying distribution of plant mating systems in our rapidly changing environments.

Unraveling these complex dynamics requires an adequate foundation of fundamental biological knowledge about plant reproduction, such as the proportion of species that can self-fertilize and how these species are distributed across the angiosperm phylogeny. To this end, we present a dataset reviewing the mating systems of 6,781 unique species, including a total of 212 families. Of these species, 1,270 are newly added from single-species studies or small reviews of under 100 species. Using these data, we explicitly address and examine potential underlying biases in the data and identify families that are over or under-sampled compared to their overall contributions to angiosperm diversity. We examined the role of existing knowledge about plant sexual systems (dioecy, monoecy, and hermaphroditism) in shaping potential sampling biases. Broadly, we revisit the question of the underlying distribution of selfing, the adequacy of our existing data, review historical change, and propose future best practices. We then extend these results to discuss bias when quantifying the frequency of selfing across angiosperms. Our results suggest that systematic study bias favoring ‘interesting’ mating systems like selfing and mixed mating impacts our understanding of mating systems in angiosperms, meaning that our understanding of this vital facet of plant life is more limited than previously thought.

### Literature Review and Scope

We conducted an extensive literature review using Web of Science, utilizing the following topic search string: “TS=((‘breeding system’ OR ‘mating system’ OR ‘self-compatib*’ OR ‘self-fertil*’ OR autogam* OR ‘auto-fertil*’ OR outcross* OR apomixis) AND plant)”. We chose these varied search terms to better capture all possible studies containing mating system information, as terminology varies significantly in plant reproductive biology (Cardoso et al. 2018). Additional filters were used to choose results from relevant journals and exclude journals that were not relevant (e.g., medical journals). The full search strings that were utilized for the review are provided in file **S1.** Journals were included from January 2017 until September 2022. This timeframe was chosen based on the existence of multiple reviews that covered existing work from the pre-2017 timeframe adequately (e.g., Goodwille et al. 2005, Moeller 2017; see **Table 1**).

**Table 1.**
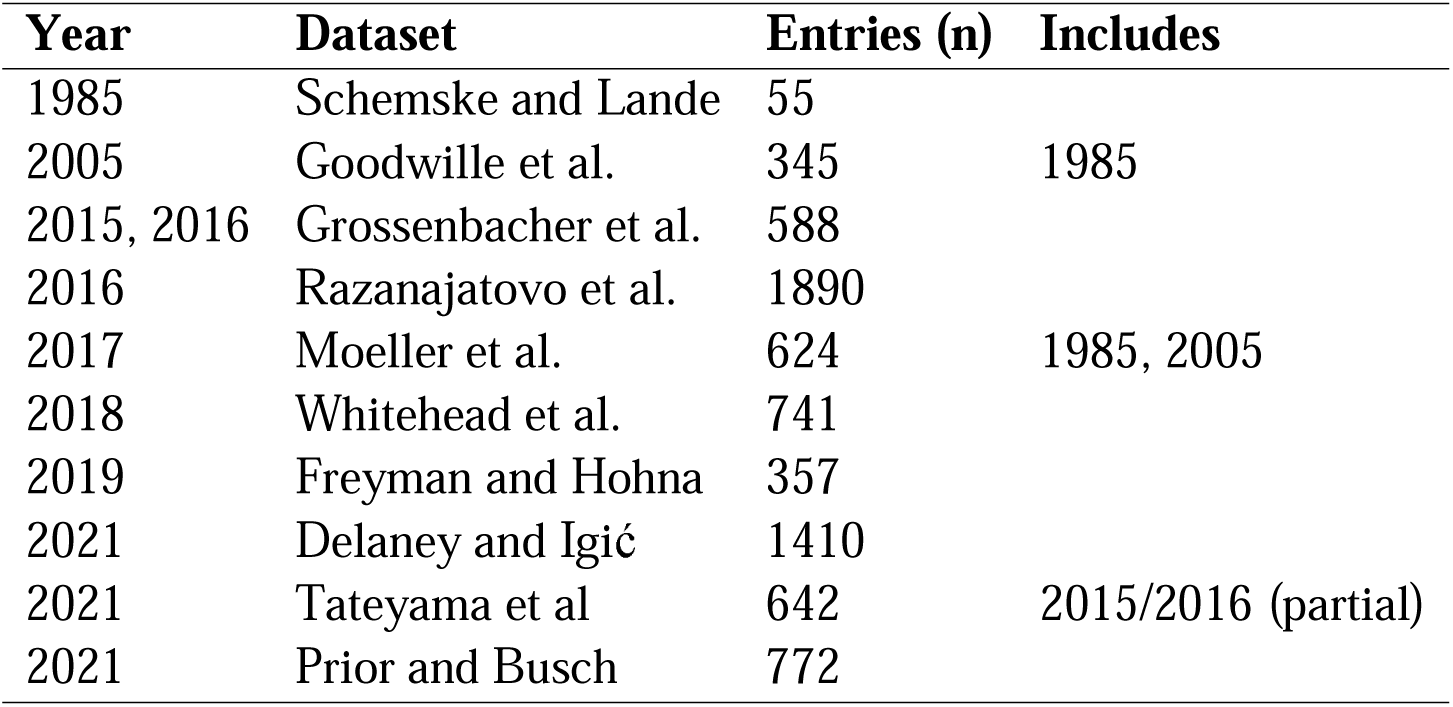
Large reviews (n>100) providing data on plant mating systems (n=11 reviews).

The filtered search produced 6,334 results. These were manually reviewed based on their abstracts for applicability. From these results, we found 1,618 studies with potentially relevant data. These studies were reviewed in full, and included if they provided, at a minimum, a binary designation of either self-compatible (SC) or self-incompatible (SI) for the focal species. In keeping with previous studies (Goodwille et al. 2005, Moeller et al. 2017), papers that reported a sexual system (i.e., apomixis) but not a selfing designation were excluded. Sexual relatives of apomictic species are generally selfing, and apomixis has been hypothesized to be correlated with selfing (Hörandl 2009, Prior and Busch 2021), so this was unlikely to impact the inclusion of outcrossing species in the dataset. Additionally, crop species were excluded, as breeding efforts and crop domestication often directly select on the ability to self (Solís Montero et al. 2021). Uncultivated crop wild relatives, which are not subjected to similar pressures, were included. Names for all species were standardized to the World Flora Online (https://www.worldfloraonline.org/), using the R package “U.Taxonstand” (R version 4.3.0, R Core Team 2023) and the WFO dataset provided by the authors (Zhang and Qian 2022). Following this review process, we included 761 studies from the 2017-2022 period. This process is fully summarized in **Figure 1**, illustrating how we processed and synthesized both existing reviews and new data into a unified dataset.

**Figure 1.**
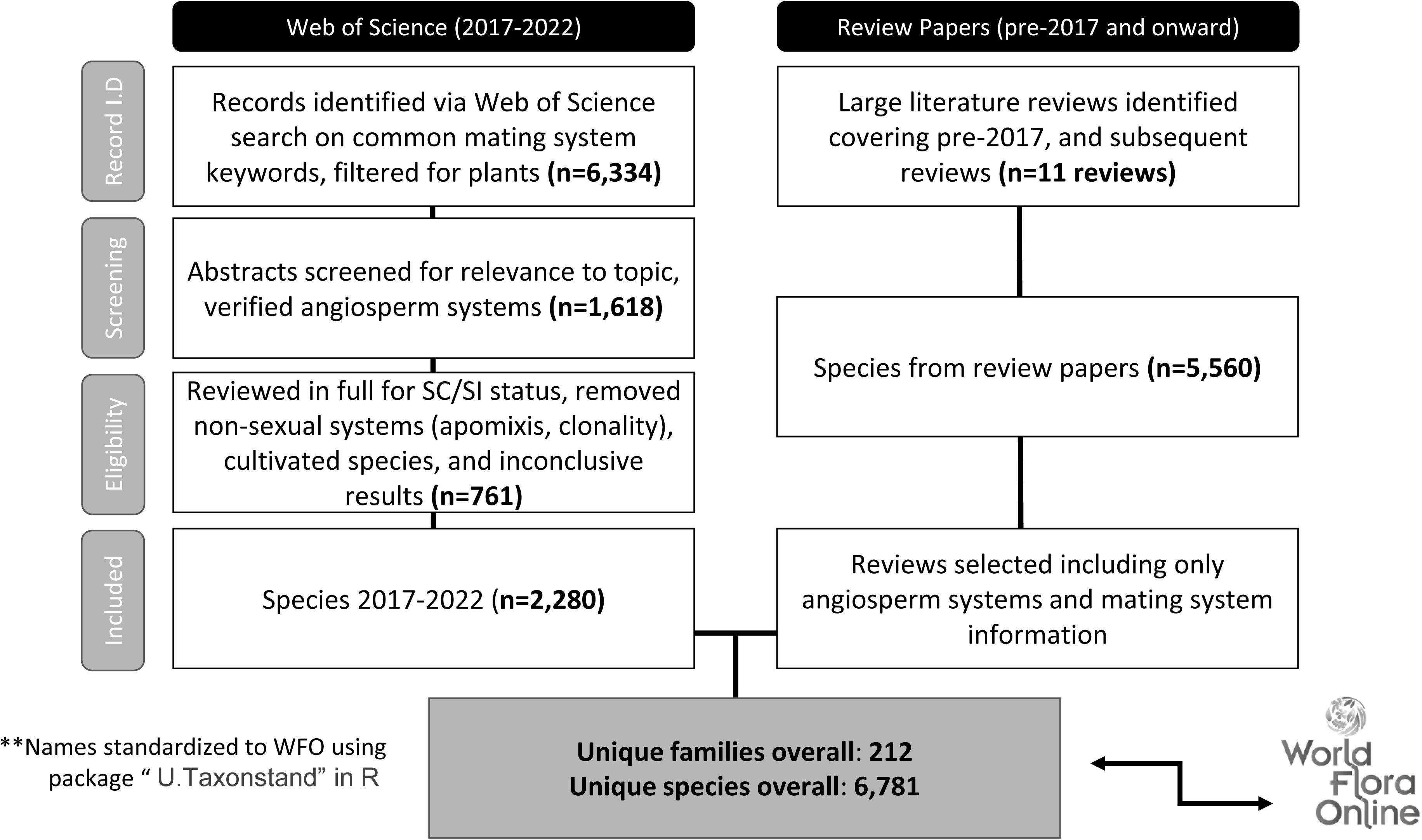
Flow chart summarizing the conducted literature review.

This literature review also included 11 large review studies (*n* > 100 species) where selfing data were also identified (Table 1). These studies were not only limited to the 2017-2022 timeframe, as they were included to represent and cover pre-2017 knowledge. These are largely synthesized in their entirety, but we excluded all non-angiosperm species (e.g., gymnosperms). While Ferrer and Good (2011) was reviewed and presented a large dataset focused on self-incompatibility/self-sterility, their work was not included in this study because the data were not accessible (i.e., numerous attempts were made to contact both authors at various time points, as well as the editorial staff at the journal in which the paper was published, but we were ultimately unsuccessful in obtaining the raw data).

Based on our synthesis of the review data (Table 1), and the 761 additional studies from 2017-2022 we newly identified from the literature, we were able to identify a total of 6,781 unique species with known mating systems. This includes 1,270 newly synthesized species. These 6,781 species draw from 212 families, or approximately half of all known families recognized by APG IV (The Angiosperm Phylogeny Group, 2016). To our knowledge, this represents the largest known single dataset on plant reproductive systems to date.

To better understand bias in sampling of mating systems, we also leveraged the greater availability of information on sexual systems. Approximately 23% of angiosperm species have a known sexual system (Wang et al. 2021). We used this as a point of reference to estimate frequencies of dioecy, monoecy, and hermaphroditism by family, based on calculating the frequency of observed species with each sexual system over the observed number of species for the family. The number of species in each family was based on Christenhusz and Byng (2016), which follows APG IV (The Angiosperm Phylogeny Group, 2016). Based on the families we identified with elevated dioecy and monoecy, we assessed representation in the mating system literature under the hypothesis that non-hermaphroditic systems would be less likely to be included in the mating system dataset. We also assessed if over-represented families had enriched hermaphroditism compared to other angiosperm families. This was done using a randomization test, in which we iteratively sampled across all families with known sexual systems (*n* = 9999 samples of 10 randomly chosen families, including over-represented families) for average percent hermaphroditism. We then compared the distribution of the randomly sampled mean hermaphroditism with that of over-represented family groups.

To assess and visualize evenness of sampling between families, we utilized a method related to rank-abundance curves from ecology. Each family in the dataset was ranked based on the total number of representative species in the dataset, and then the number of species represented was normalized to the family with the highest number of species. We also examined how this changed over time using the reviews of Schemske and Lande (1985), Goodwille et al. (2005), and Moeller (2017).

## Results and Discussion

### Few families have well-studied mating systems proportional to overall diversity within families

Our goal was to better understand the state of knowledge of plant reproductive systems across all angiosperms. To cover the pre-2017 period, we synthesized major reviews of selfing, starting with Schemske and Lande (1985). To cover the 2017-2022 period, we reviewed 761 papers comprising 6,781 unique species. This species list has at least one representative from 217 families, or approximately half (52.2%) of all known angiosperm families (Christenhusz and Byng 2016). To the best of our knowledge, this is the largest such dataset presented to date, both with respect to the number of species, and the total number of families represented.

From our dataset, 63% of species were SC and thus selfed or had the potential to self at least some of the time. This includes both obligate selfing species and mixed mating species. An additional 37% were SI and hence obligately outcrossing. Species with multiple conflicting designations (i.e., studies indicating both SC and SI for the same species) were excluded. This decision was based on similar practices in other mating system studies (Moeller et al. 2017, Prior and Busch 2021). These species with conflicting designations comprised 7.5% of the dataset. In contrast to our results, Goodwille et al. (2005) reported 42% of their sample as mixed mating. While our proportion of SC species was larger, our sample includes SC species that would have been excluded from the mixed mating sample of Goodwillie et al. (2005) based on their definition of mixed mating. Igić and Kohn’s (2006) discussion of potential biases in mating system studies proposed a corrected 48.7% frequency of SI based on their estimation of under-sampling of obligate outcrossing species.

While our review synthesized a large number of studies, known mating system data for selfing remains limited compared to overall angiosperm diversity. Additionally, sampling is highly uneven across family groups. Thus, despite the size of our dataset, many families have a limited coverage of taxonomically described species. For example, out of the 212 families included here, only 51 families had more than 5% of their total species represented, while only 34 families had more than 10% of their total species represented. Additionally, only 14 of these 34 families had more than 100 species covered, suggesting limited understanding of larger family groups. To illustrate these trends, the coverage for larger (*n* > 500 sp.) families is summarized in **Table 2**. This shows how many large families have limited representation, and a small number of families are significantly better studied than others.

**Table 2.** Within-family coverage obtained in our review of plant mating systems, for families of 500 species or more with 3% or greater coverage in our dataset.

| Family | Species (n) | Coverage (%) |
| --- | --- | --- |
| Onagraceae | 656 | 48.63 |
| Solanaceae | 2600 | 20.04 |
| Brassicaceae | 3628 | 9.40 |
| Lythraceae | 620 | 9.19 |
| Geraniaceae | 830 | 8.43 |
| Fabaceae | 19500 | 6.96 |
| Bignoniaceae | 870 | 4.37 |
| Bromeliaceae | 3475 | 4.35 |
| Balsaminaceae | 1000 | 3.90 |
| Proteaceae | 1660 | 3.80 |
| Asteraceae | 24700 | 3.62 |
| Plantaginaceae | 1900 | 3.53 |
| Dipterocarpaceae | 695 | 3.45 |
| Cactaceae | 1750 | 3.37 |

Indeed, a disproportionate amount of the overall dataset comes from a limited number of well-covered families, which differs drastically from these families’ contribution to overall angiosperm diversity. If species were randomly sampled across the angiosperm phylogeny, these percent coverages should more closely match. For example, species classified within Solanaceae represent 7.7% of our data, but with its approximately 2,600 species covered at 20.04% in our dataset, this family represents only 0.9% of total estimated angiosperm species. Similarly, Asteraceae, Fabaceae, Brassicaceae, Onagraceae, and Bromeliaceae are all overrepresented in our dataset compared to their expected proportions based on family size if random sampling was used as the criteria to select species for inclusion in our dataset. Some of the largest families, moreover, are notably underrepresented: Orchidaceae, Rubiaceae, and Poaceae (**Figure 2**).

**Figure 2.**
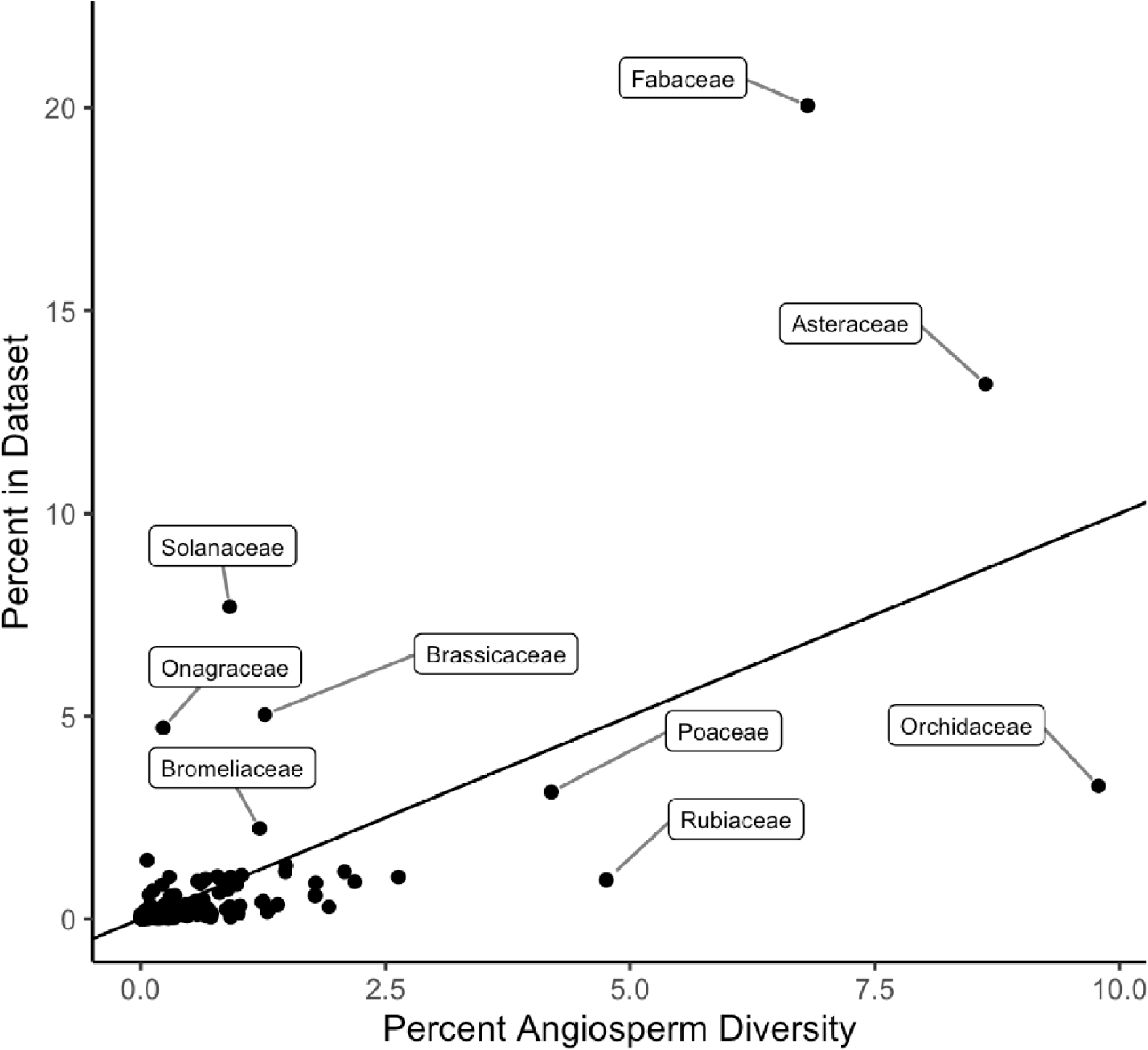
Families represented in our mating system dataset plotted by the percent of angiosperm diversity they represent (x-axis) and the percent of our dataset they represent (y-axis). Points falling above the 1:1 line are over-represented, points falling below it are under-represented.

It is important to note that our dataset represents a very small portion of the overall diversity of flowering plants. There are 295,383 known plant species recognized in APG IV (Christenhusz and Byng 2016), meaning that out of all species, only approximately 2% have a documented mating system in our dataset. For comparison, plant sexual systems (i.e., monoecious, dioecious, or hermaphrodite) are known for at least 23% of all species in APG IV (Wang et al. 2021). While determining the mating system of a species can be more challenging than determining the sexual system, this is nonetheless an example of a related floral trait for which greatly more information is available.

### Existing data arises disproportionally from a small number of families with labile mating systems

A small number of families comprised the majority of existing plant mating system data. The top 10 best-covered families in our review contained over half of the total species (50.90%). In contrast, these 10 families contribute only 1.2% of overall angiosperm diversity. Similarly, if we look at the top 10 families in our review by raw number of species with known mating systems (instead of percent coverage), these top 10 families represent 64.7% of the total species in the review, but still only 33.51% of total angiosperm biodiversity. We calculated rank-abundance plots for Schemske and Lande (1985), Goodwille et al. (2005), Moeller et al. (2017), and our study to quantify trends in sampling dominance over time (**Figure 3**). For comparison, these are shown alongside the expected angiosperm-wide rank-abundance curve using current estimates of angiosperm diversity and taxonomy. As time progresses, the rank-abundance curves for sampled species more closely match the expectation given our current taxonomic knowledge. However, as shown in **Figure 4**, the most-abundant families making up the steep top of the abundance curves are not the same families as expected. If sampling was truly random across angiosperms, Orchidaceae, Asteraceae, Fabaceae, Rubiaceae, and Poaceae – the most abundant families in our current understanding of angiosperm diversity – would form the top of the curve in all instances, especially those later in time. This is clearly not the case. Furthermore, the contributions of these families to our collective understanding of plant mating systems varies over time and across studies to such an extent that it is questionable whether summaries of existing data are unbiased.

**Figure 3.**
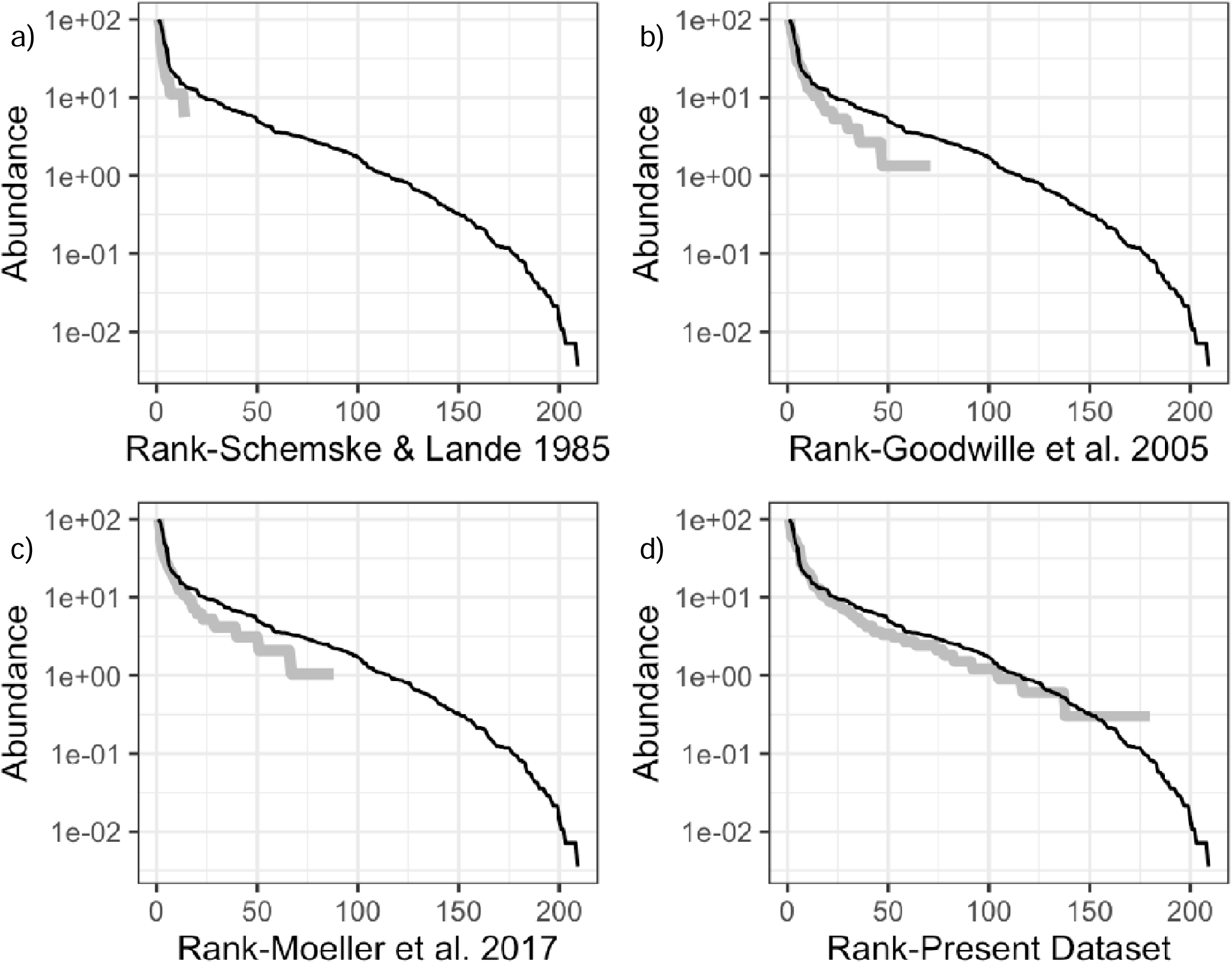
Log-scale rank-abundance plots. Rank-abundance curves are shown for three cumulative studies: a) Schemske and Lande (1985), b) Goodwille et al. (2005), and c) Moeller et al. (2017), as well as d) the present study.

**Figure 4.**
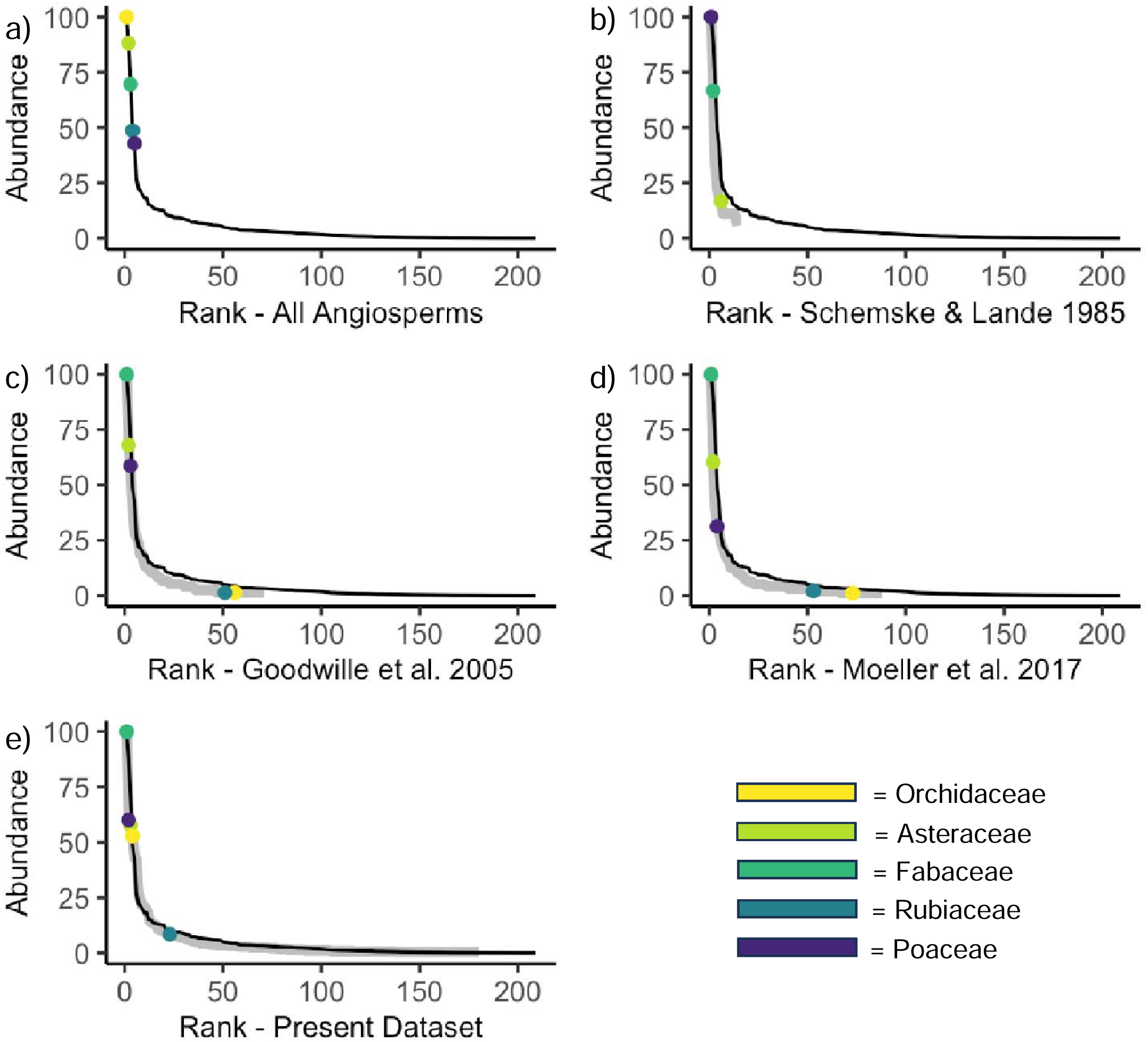
Rank-abundance curves for b) Schemske and Lande (1985), c) Goodwille et al. (2005), d) Moeller et al. (2017), and e) the present study, showing the relative positions of the most abundant angiosperm families in their respective datasets. The top left panel (a) shows the expected abundance based on relative angiosperm family diversity. (Note that Orchidaceae and Rubiaceae do not appear in Schemske and Lande (1985) and thus are not plotted on the curve.)

Some of the unevenness in sampling across families likely arises from studies focused on using PCMs to describe evolutionary transitions to and from selfing. It is notable that several of the best-studied families (Table 2) have also been the focus of PCM studies (Solanaceae, Goldberg et al. 2010; Onagraceae, Freyman and Hoha 2019; Fabaceae, Igić and Delaney 2022). One can also imagine this interaction in the opposite way; these families were chosen for PCM study due to the greater data availability. Regardless of the direction of this interaction, the inevitable result is that our understanding of the evolution, ecological characteristics, and frequency of selfing arises disproportionately from a limited number of families that may not be representative of angiosperms overall.

Choosing families with labile mating systems is appropriate when using PCMs to estimate rates of evolutionary transitions to and from selfing, as well as the correlations of these transitions with overall diversification rates. Due to better availability of trait data at tree tips, statistical power for parameter estimation using PCMs is maximized for these families (Harmon 2018). In Onagraceae, for example, autogamy has long been known to evolve easily, and has evolved at least 150 times independently (Raven 1979). Similarly, Solanaceae is known to have intermediate levels of self-incompatibility and frequent mating system transitions (Goldberg et al. 2010, 2012). The only other family with similarly high coverage with respect to mating system data is Phrymaceae, which contains the genus *Mimulus* (previously classified in the now defunct Scrophulariaceae), a frequent model system for studies of selfing and ecological genomic studies (Wu et al. 2008). Perhaps unsurprisingly, in our dataset, all representatives of Phrymaceae come from the genus *Mimulus.* Studies of selfing in *Mimulus* could explain why more mating system information exists for this particular family, as it has become a model system for many different questions in ecology and evolution. In a similar way, the field of genetics is grappling with model organism choice in an era of increased availability of genomic data, with some arguing that existing model organisms are insufficient to capture the diversity and complexity of the systems they are supposed to represent (Bertile et al. 2023). The study of plant reproductive systems likely faces a similar problem: families that are used as model systems, and thus contribute disproportionately to available knowledge on mating systems, are not representative of angiosperm diversity as a whole.

### Families with an elevated frequency of dioecious or monoecious species are less likely to be well-studied

Considering the uneven sampling shown above, we also utilized more-available information about plant sexual systems (categorized here as monoecy, dioecy, and hermaphroditism) to examine potential biases in mating system studies. This is sensible given the clear biological links between sexual systems and mating systems. For instance, if a bias existed against families with a significant number of dioecious species, this would suggest a bias against obligate-outcrossing species. Similarly, self-fertilization in monoecious species could be more challenging than in their hermaphroditic counterparts, and under-sampling of families with known high levels of monoecy could also suggest bias based on preexisting knowledge about sexual system and morphology.

For dioecy, the relationship between sexual system and mating system is straightforward: dioecious plants are morphologically and definitionally unable to self. Dioecy is comparatively uncommon in sexual systems (Renner and Ricklefs, 1995), but some families have considerably enriched dioecy (*n* = 15 families with > 200 species). These families had between 10%-49% dioecious species (Wang et al. 2021). From these highly dioecious families, all 15 rank in the bottom 50% of families for mating system coverage in our dataset, and all but two rank in the bottom 30%. Additionally, one third of these families (Smilacaceae, Cucurbitaceae, Vitaceae, Putranjivaceae, and Menispermaceae) did not have a single species with a known mating system listed in any published study (**Figure 5**). This is especially surprising as Cucurbitaceae and Vitaceae are families with economically important species.

**Figure 5.**
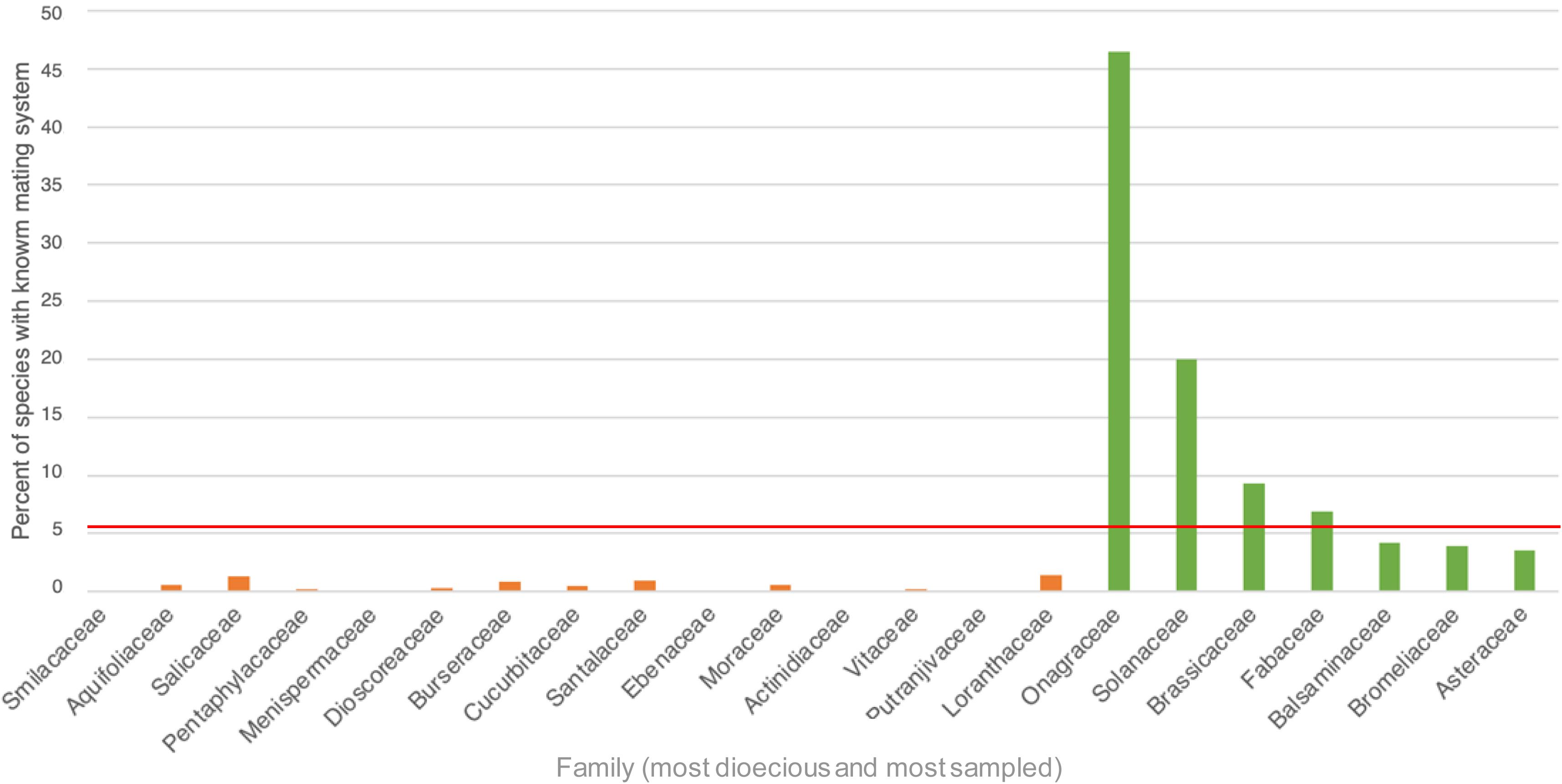
The percent coverage for these families is shown alongside six of the best-covered families for scale. The red line represents the average percent coverage across all families, at 6.95% coverage.

The relationship between the mating system and sexual system for monecious species is more complex. While monoecy is frequently hypothesized to be an adaptation to limit selfing, empirical evidence is limited: Bertin (1993) theorized if monoecy was an adaptation to prevent selfing, SI would be uncommon in species exhibiting this sexual system. Unexpectedly, Bertin (1993) found monoecy to be independent of the mating system. However, for our data, highly monoecious families (defined as families with >5% monoecy) were also underrepresented in our dataset relative to their expectation based on overall angiosperm diversity, and we found that the 15 families with the greatest levels of known monoecy fall well below the mean level of coverage in the plant mating systems data (Table 4). As a corollary to this, families with high levels of dioecious and monoecious species typically have very limited hermaphroditic species. While an incomplete sample of species have a known sexual system, most families that are exhibiting high dioecy or monoecy also have very limited or no hermaphroditism. The percent hermaphroditism for each family is shown in **Tables 3 and 4**, which shows that underrepresentation of monoecy and dioecy likely also yielded an enrichment in hermaphroditism.

**Table 3.**
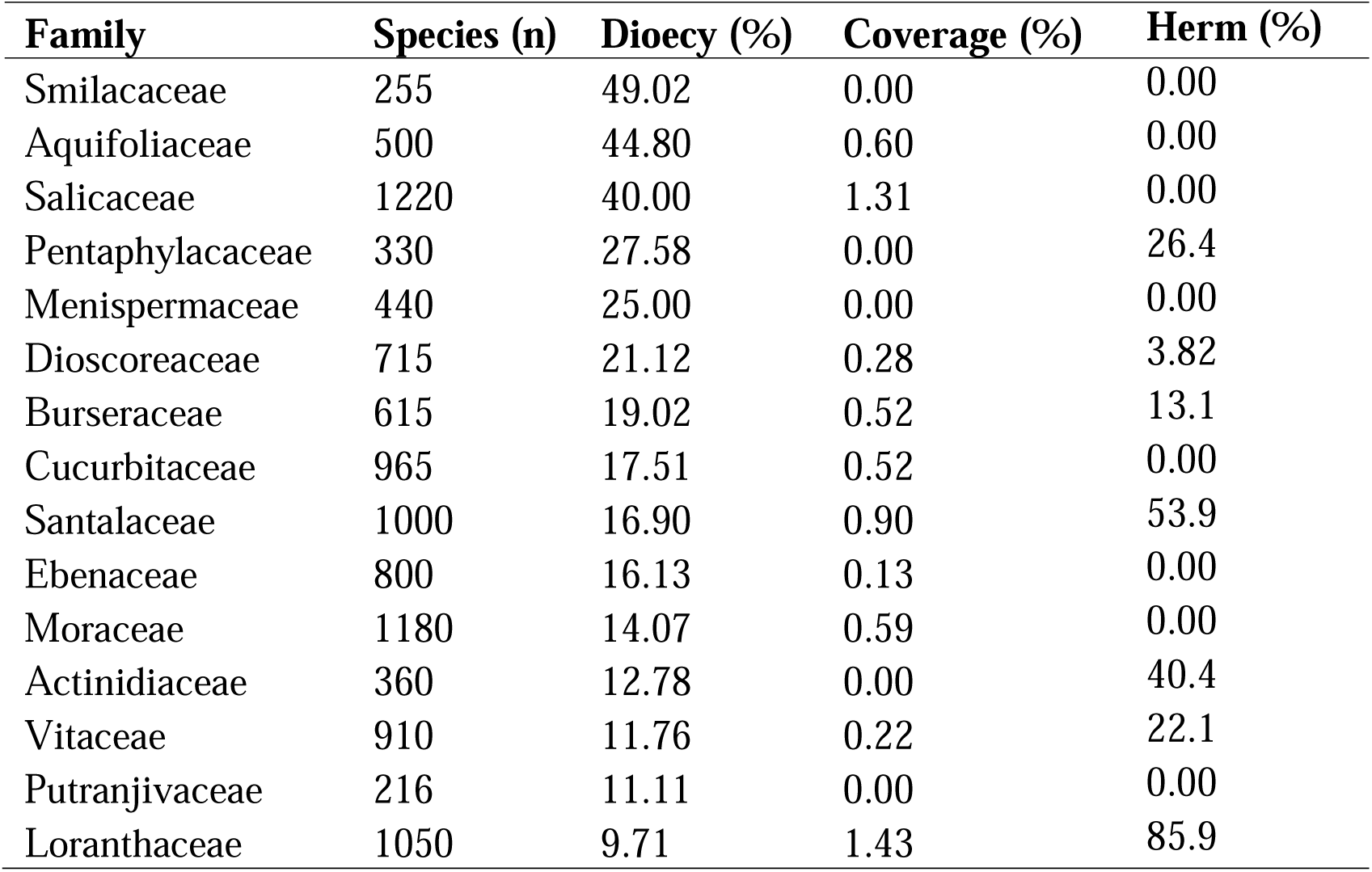
Families with enriched dioecy based on Wang et al. (2021), by percent of known dioecious species relative to overall family size. Percent of family known to be hermaphroditic is also shown.

**Table 4.**
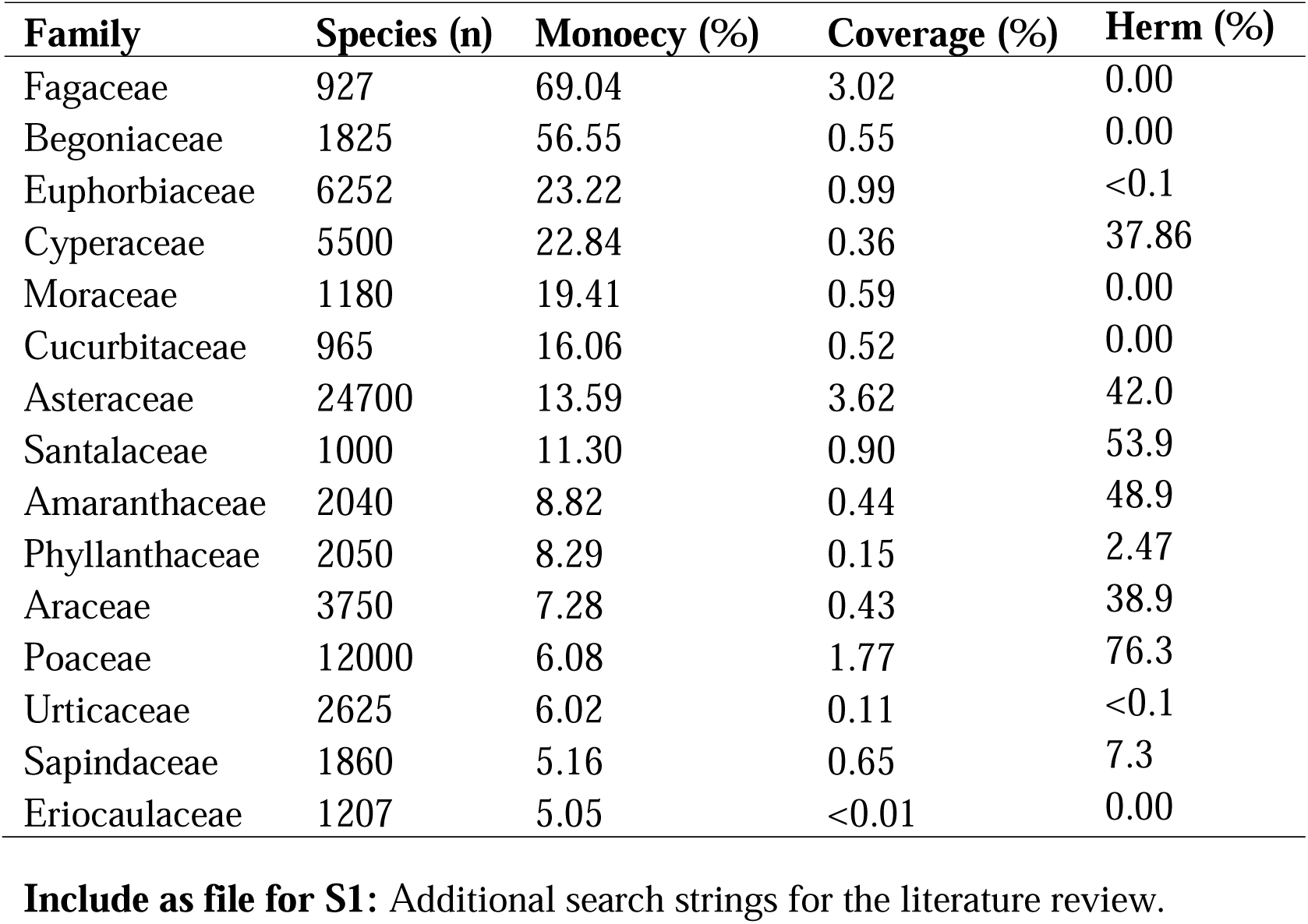
Families with enriched monoecy based on Wang et al. (2021), by percent of known monoecious species relative to overall family size. Percent of family known to be hermaphroditic is also shown.

We examined the families that were over-represented with respect to selfing information (**Figure 2**) and found these families to be enriched for hermaphroditism (**Figure 6**). The mean hermaphroditism for families in Wang et al. (2021) was 72%, and the mean for Solanaceae, Asteraceae, Fabaceae, Brassicaceae, Onagraceae, and Bromeliaceae was 87%. To better quantify this deviation from the expected mean, we conducted an ad-hoc randomization test. We iterated 9999 random draws of six families from the 330 families with known sexual systems in Wang et al. (2021) and generated a frequency distribution of mean hermaphroditism. Our mean hermaphroditism (72%) for over-represented families can be seen to fall to the right tail of the distribution (**Figure 6**). This suggests that not only are dioecious and monoecious species under-sampled, but hermaphroditic species are over-sampled. This is consistent with a general bias against outcrossing species, which leads to a positive bias for selfing species. Thus, existing data, even when expanded to sexual systems, likely overestimates the frequency of selfing species.

**Figure 6.**
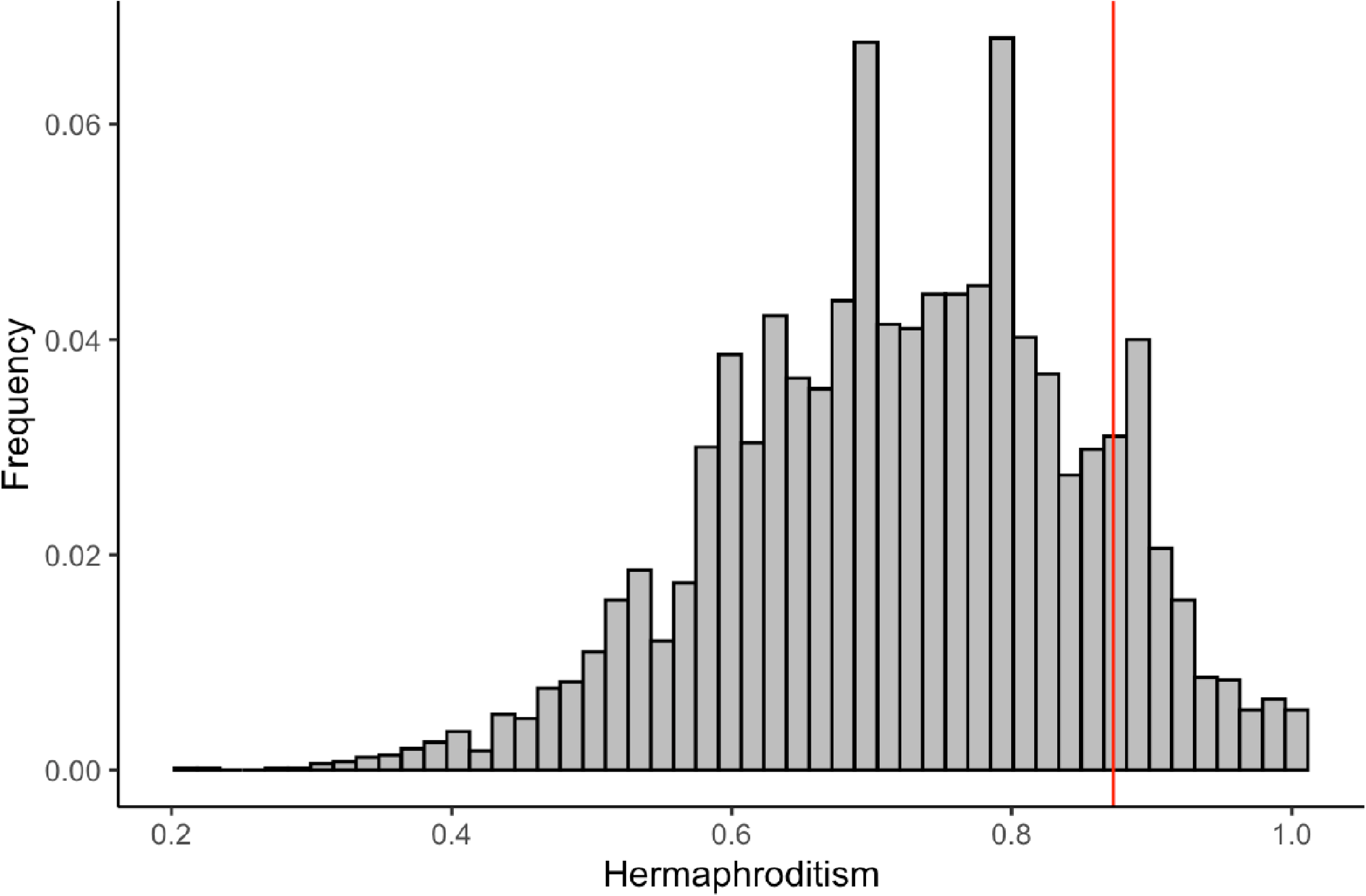
The frequency distribution of mean hermaphroditism in families resulting from a randomization test. The read line indicates the mean for families over-represented in our dataset, at 0.8725.

Data about plant sexual systems are also incomplete. However, persistent undersampling of dioecious and monoecious families points to a larger pattern of under-representation of outcrossing in mating and sexual system studies. These results suggest that species known to have limited capacity for selfing due to their anatomy are less likely to be included in databases about plant mating systems. Furthermore, as selfing is correlated with a wide variety of other traits – the so-called “selfing syndrome” (Darwin, 1879, Ornduff 1969) – this could directly impact our understanding of other traits in angiosperms using the same resources as presented here.

Based on our survey of mating system literature, we initially produced an estimate of 62.5% SC and 38.5% SI. However, in light of the biases we discuss above, we must consider that these estimates over-estimate SC, and under-estimate SI. Based on our synthesis of sexual system data, we could leverage this information to make some informed choices when accounting for bias in mating system data. Based on Wang et al. (2021), dioecy accounts for about 8% of total plant species, monoecy 16%, and hermaphroditism 76%. While the family-level under and oversampling suggests a larger issue of focus on SC and mixed mating, a simple initial correction could be to correct for limited dioecy. For species we studied with both mating system and sexual system data (n=2217), our sample has about 2% dioecy, 13% monoecy, and 85% hermaphroditism. If we were to adjust for under-representation of SI contributions from dioecy and monoecy (assuming dioecious species are always SI, and 63% of monoecy species are SI, based on our data), we would have a new estimate of 58% SC and 42% SI. But as noted previously, the patterns of sampling bias that produced the observed deficiency of dioecy and monoecy in mating system data could also impact sampling in hermaphroditic species. Thus it seems entirely plausible that our existing estimates remain SC-enriched.

### Conclusions and Future Directions

By assembling existing knowledge of selfing in an updated, harmonized, and publicly available dataset, we hope to promote further research in the rich field of plant reproductive systems. We especially hope to encourage studies that address the underlying distribution of selfing in angiosperms, and selfing-focused papers broadly to consider the adequacy of existing data. We also hope to promote additional investigation into as-of-yet understudied groups, such as understudied families with dioecious and monoecious species, or families that have been shown to be especially under-represented compared to their species diversity.

As we have shown, obligate outcrossing species may be systematically underrepresented, which was previously shown by Igić and Kohn (2006). We have additionally shown that this bias is likely informed by prior knowledge about families, leading to “interesting” mating systems like selfing and mixed mating being over-studied. Additionally, a small number of families represent a large amount of existing knowledge about plant mating systems. Some of these families may contain a higher-than-average number of selfing species based on enriched hermaphroditism. In order to address this problem, we need not only *more* data, but thoughtfully collected data. For example, data that is collected randomly or data focusing on improving knowledge of understudied groups would be especially useful at this juncture. As illustrated by our work, it will also be vital that we utilize a more holistic view of plant reproduction. Sexual systems and other mating-system associated traits that are more available can be leveraged to help us generate reasonable estimates of selfing or to assess the data that we *do* have. For example, authors studying mating systems within specific family groups would be well-advised to examine family-level data about frequency of SI, SC, and different sexual systems within their family, and to consider critically the adequacy of that data and potential interplay between mating and sexual systems.

Our results indicate that we understand less about the underlying frequency of selfing across angiosperms than was previously thought. Self-fertilization presents a potential route for reproductive assurance for plants in challenging and unstable environments (Baker 1955, Busch and Delph 2012), but can have significant determinants as well. In an era of unprecedented anthropogenic disturbance, it thus seems plausible that an individual plant’s reproductive strategy could become a stronger predictor of its fitness than it has been in the past. Reproductive systems are known to be labile in many large families and across populations within species (Whitehead et al. 2018).

It could be imagined that under anthropogenic climate change, selfing could be the way of the future as reproductive assurance is needed for short-term survival of some species. However, in the long term, the known deleterious consequences of inbreeding could limit fitness or lead to extinction in selfing species. This could represent an as-of-yet unforeseen cost of anthropogenic change on angiosperm diversity. This could even illuminate a new feedback loop of biodiversity loss in plants, a foundational piece of all terrestrial ecosystems. This illustrates the importance of understanding the prevalence of outcrossing, selfing and mixed mating across angiosperms.

## Acknowledgments

The authors thank Michael S. Rosenberg, Bret M. Boyd, and Rodney J. Dyer for many discussions of this work, as well as the Integrative Life Sciences Doctoral Program at Virginia Commonwealth University for ongoing support of this project.

All authors contributed to study design and conception. EM Meyer conducted the literature review, analysis, and drafted the manuscript. All authors contributed to following subsequent revision.

## Notes

### Competing Interest Statement

The authors have declared no competing interest.

### Summary of Updates

Abstract updated to a structured abstract; copy-edits; small figure revisions for clarity.

