## Supplemental Text for "The evolutionary dynamics of plant mating systems: how bias for studying ‘interesting’ plant reproductive systems could backfire"

**Include as file for S1:**

Additional search string used to filter for journals relevant to the literature review and timeframe: “and **Agronomy** or **Plant Sciences** or **Ecology** or **Genetics Heredity** or **Evolutionary Biology** or **Horticulture** or **Multidisciplinary Sciences** or **Environmental Sciences** or **Biology** or **Biodiversity Conservation**or **Forestry** or **Reproductive Biology** or **Developmental Biology** or **Environmental Studies** or **History Philosophy Of Science** or **Anatomy Morphology** (Web of Science Categories) and **2017** or **2018** or **2019** or **2020** or **2021** or **2022** (Publication Years)”.
